# Human milk oligosaccharides reduce murine group B *Streptococcus* vaginal colonization with minimal impact on the vaginal microbiota

**DOI:** 10.1101/2021.10.20.465155

**Authors:** Marlyd E. Mejia, Samantha Ottinger, Alison Vrbanac, Priyanka Babu, Jacob Zulk, David Moorshead, Lars Bode, Victor Nizet, Kathryn A. Patras

## Abstract

Group B *Streptococcus* (GBS) colonizes the vaginal mucosa of a significant percentage of healthy women and is a leading cause of neonatal bacterial infections. Currently, pregnant women are screened in the last month of pregnancy and GBS-positive women are given antibiotics during parturition to prevent bacterial transmission to the neonate. Recently, human milk oligosaccharides (HMOs) isolated from breastmilk were found to inhibit GBS growth and biofilm formation *in vitro*, and women that make certain HMOs are less likely to be vaginally colonized with GBS. Using *in vitro* human vaginal epithelial cells and a murine vaginal colonization model, we tested the impact of HMO treatment on GBS burdens and the composition of the endogenous microbiota by 16S rRNA amplicon sequencing. HMO treatment reduced GBS vaginal burdens *in vivo* with minimal alterations to the vaginal microbiota. HMOs displayed potent inhibitory activity against GBS *in vitro*, but HMO pretreatment did not alter adherence of GBS or the probiotic *Lactobacillus rhamnosus* to human vaginal epithelial cells. Additionally, disruption of a putative GBS glycosyltransferase (Δ*san*_0913) rendered the bacterium largely resistant to HMO inhibition *in vitro* and *in vivo* but did not compromise its adherence, colonization, or biofilm formation in the absence of HMOs. We conclude that HMOs are a promising therapeutic bioactive to limit GBS vaginal colonization with minimal impacts on the vaginal microenvironment.

**IMPORTANCE:** During pregnancy, GBS ascension into the uterus can cause fetal infection or preterm birth. Additionally, GBS exposure during labor creates a risk of serious disease in the vulnerable newborn and mother postpartum. Current recommended prophylaxis consists of administering broad-spectrum antibiotics to GBS-positive mothers during labor. Although antibiotics have significantly reduced GBS neonatal disease, there are several unintended consequences including altered neonatal gut bacteria and increased risk for other types of infection. Innovative preventions displaying more targeted antimicrobial activity, while leaving the maternal microbiota intact, are thus appealing. Using a mouse model, we found that human milk oligosaccharides (HMOs) reduce GBS burdens without perturbing the vaginal microbiota. We conclude that HMOs are a promising alternative to antibiotics to reduce GBS neonatal disease.

## INTRODUCTION

Group B *Streptococcu*s (GBS or *Streptococcus agalactiae*) is a Gram-positive bacterium that colonizes the gastrointestinal and vaginal tracts of ~18% of pregnant women globally (1), exposing more than 20 million infants to GBS at, or prior to, delivery (2). The majority of children born to GBS-positive women themselves become colonized without symptoms (3); however, a subset of these infants (>300,000 annually) develop invasive GBS infections accounting for upwards of 100,000 infant deaths each year around the globe (2). Additionally, 57,000 annual stillbirths are attributed to GBS infections (2), yet this may be an underestimate as this pathogen is also the most frequently cultured bacterium in mid-gestation spontaneous abortions (4). Because maternal GBS colonization is a risk factor for neonatal infections, universal screening in late pregnancy and intrapartum antibiotic prophylaxis (IAP) to GBS-positive or at-risk mothers is the current standard of care in many countries. These preventative measures have decreased, but not eradicated, GBS early-onset disease (5). However, this early antibiotic exposure disrupts the infant microbiota and the potential adverse consequences of this perturbation are not fully established (6-10).

Breastfeeding has long been associated with improved infant health, reduced risk of infectious disease, and accelerated immune and microbial maturation within the gut (11-13). Human milk oligosaccharides (HMOs), the third most abundant component of breastmilk, are a group of structurally complex, unconjugated glycans that are recalcitrant to host digestive enzymes. HMOs provide nutritional advantage for beneficial microbes in the infant gut and drive immune maturation at the gut epithelium (13-16). Moreover, HMOs may protect against neonatal pathogens by acting as soluble “decoy” receptors for enteric pathogens (17, 18), through neutralization of bacterial toxins (19, 20), or via direct antimicrobial activity including against GBS (21-24). Although the mechanism of HMO-mediated GBS inhibition is not known, GBS expression of a putative glycosyltransferase (locus *san*_0913) is necessary for inhibitory activity (21), and HMO exposure lowers GBS sensitivity to antibiotics including vancomycin, erythromycin and trimethoprim (21, 25, 26). Additional support for HMO-mediated anti-GBS activity stems from clinical observations that mothers who produce a functional variant of the fucosyltransferase enzyme *FUT3*, which attaches fucose in an α1–3 or α1–4 linkage to form certain HMOs, are less likely to be vaginally colonized by GBS (27).

We hypothesized that HMOs may reduce GBS vaginal colonization *in vivo* either through direct antimicrobial activity, or through indirect activity on the vaginal epithelium and/or vaginal microbiota. Here, we test this hypothesis using a murine model of GBS vaginal colonization and pooled HMOs (pHMOs) isolated from human breastmilk. We further assess the impact of pHMOs on bacterial attachment to human vaginal epithelial cells and phenotypically characterize a GBS strain that is resistant to HMO inhibitory actively (21). Combined, our findings support the continued exploration of HMOs as a therapeutic strategy for GBS in pregnancy and the neonatal period.

## RESULTS

### Topical pHMO treatment reduces GBS vaginal burdens *in vivo*

To determine the effect of HMOs on GBS vaginal colonization *in vivo*, wild-type C57BL/6J female mice were vaginally inoculated with GBS COH1, a serotype III ST17 neonatal sepsis clinical isolate (28). Mice were treated with pHMOs (1 mg/dose) 2 h prior to GBS inoculation, and on the following two consecutive days. Lacto-N-tetraose (LNT), a commercially produced HMO that inhibits GBS growth *in vitro* (21) was included as a treatment condition to test the efficacy of a single HMO. Vaginal swabs were collected prior to pHMO treatment on day 0, 1, and 2, as well as day 3 and 6 post-inoculation (**Fig. 1A**). Treatment with pHMOs significantly reduced GBS vaginal burdens on day 1 (*P* = 0.023) and 2 (*P* = 0.009) during active treatment, but these differences were resolved at day 3 and 6 after pHMO treatment had stopped (**Fig. 1B**). No differences between LNT and mock-treated groups were observed at any time point. Additionally, endogenous vaginal *Enterococcus* spp. were distinguished on the *Streptococcus* selective media, but no differences between treatment groups were detected (**Fig. 1C**).

**Figure 1.**
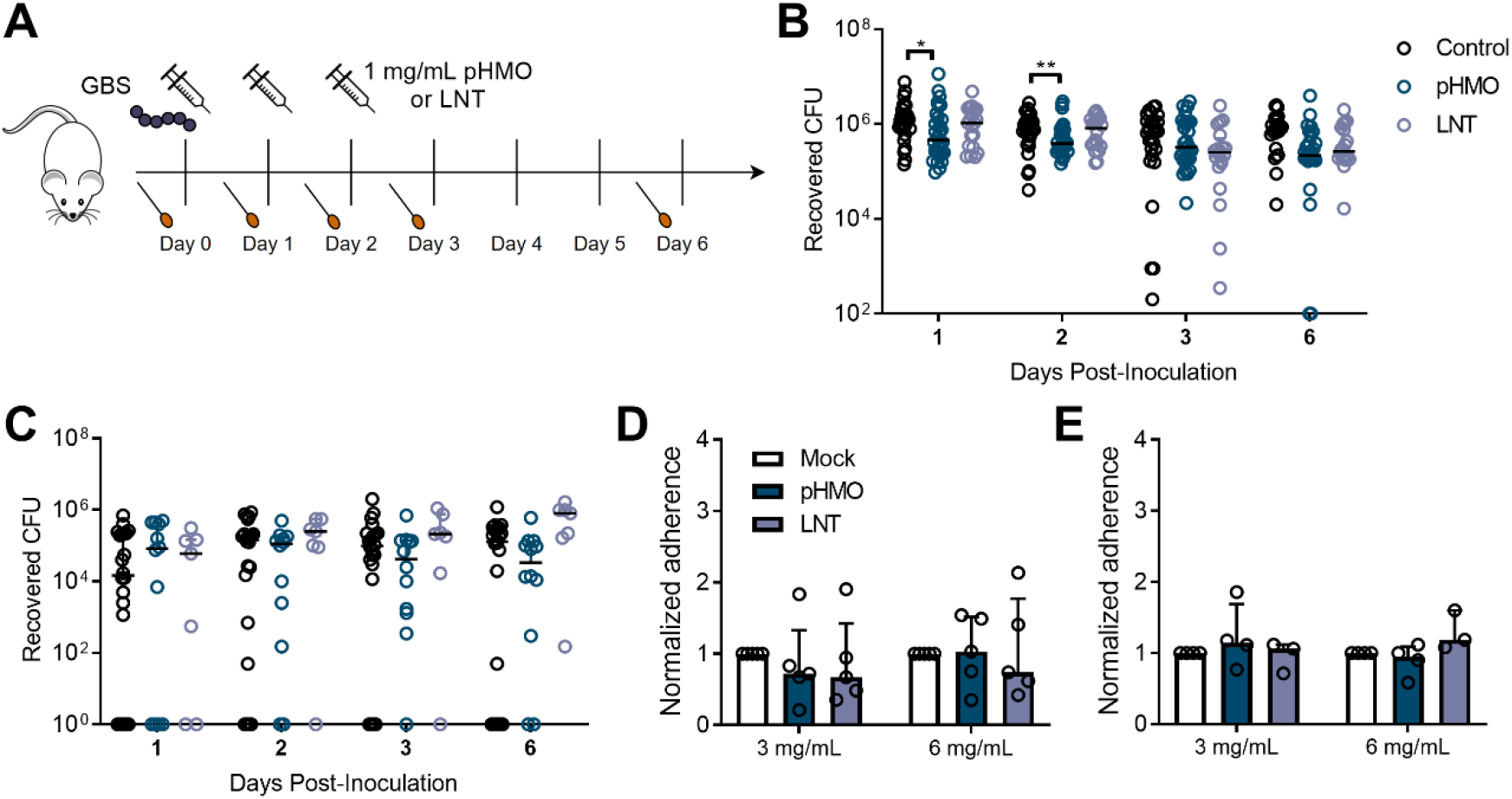
Treatment with pHMOs, but not specific HMO LNT, reduce GBS vaginal burdens in mice and do not impact adherence to human vaginal epithelial cells. (A) Experimental timeline for the GBS colonization model. Baseline vaginal swabs were collected on Day 0 prior to GBS inoculation with 1 × 10^7^ CFU of GBS COH1. Mice were treated with 1 mg pHMOs or lacto-N-tetraose (LNT) two hours post-infection, and on the two subsequent days. Mice were swabbed prior to daily treatment with HMOs, as well as one and four days after the last HMO treatment. Mouse and syringe images are available open source through pixabay. (B) GBS burdens recovered from mouse vaginal swabs over the 6-day time course (*n* = 20-30/group). (C) *Enterococcus* spp. burdens recovered from mouse vaginal swabs over the 6-day time course. Mice that did not culture *Enterococcus* at any time point were excluded (*n* = 7-22/group). Adherence of GBS COH1 (D) or *Lactobacillus rhamnosus* GG (E) to VK2 cells pretreated with pHMOs or LNT for 18 h. Adherence was normalized to mock treated controls. Symbols represent individual mice (B, C), or the means of 4-5 independent experimental replicates (D,E), with lines representing median and interquartile range. Data were analyzed by Kruskal Wallis with Dunn’s multiple comparisons test (B,C) or two-way ANOVA with Dunnett’s multiple comparisons test (D,E). ** *P* < 0.01, * *P* < 0.05. All other comparisons are not significant.

### Vaginal epithelial HMO exposure does not impact adherence of GBS or probiotic *Lactobacillus*

Because HMOs can reduce adherence of pathogens (29-31) and promote adherence of beneficial bacteria to the host epithelium (32), we tested the impact of epithelial HMO pretreatment on adherence of GBS or the probiotic *Lactobacillus rhamnosus* GG to human vaginal epithelial (VK2) cells. We observed no effect of pHMO or LNT pretreatment on GBS adherence to VK2 cells at two different concentrations (**Fig. 1D**), nor did HMO pretreatment alter *L. rhamnosus* adherence to VK2 cells (**Fig. 1E**).

### HMO resistance conferred by disruption of *san*_0913 does not alter GBS biofilm formation, adherence, susceptibility to antibiotics, or *in vivo* colonization in the absence of HMOs

Although the exact mechanism of HMO anti-GBS activity has yet to be established, increased GBS sensitivity to intracellular targeting antibiotics and enhanced cell membrane permeability occur following HMO exposure (21, 25, 26). Additionally, HMO exposure perturbs multiple GBS metabolic pathways including those related to linoleic acid, sphingolipid, glycerophospholipid, and pyrimidine metabolism (26). A transposon mutant library screen identified the *gbs0738* gene (locus *san*_0913 or GBSCOH1_RS04065 in COH1), a putative glycosyltransferase family 8 protein, as essential for GBS susceptibility to HMOs over a 7 h time course (21). Using a targeted insertional mutant of *san*_0913 (COH1 Δ*san*_0913) (21), we assessed the growth of WT COH1 and Δ*san*_0913 in the presence of 0-20 mg/mL pHMOs over 18 h. We found that growth of COH1 was significantly inhibited at all pHMO concentrations tested compared to the mock control (**Fig. 2A,B**). Concentrations of 20 mg/mL and 10 mg/mL pHMO inhibited growth of Δ*san*_0913 but to a lesser degree than seen with wild-type COH1 (**Fig. 2A,B**). To determine whether *san*_0913 disruption altered GBS characteristics associated with colonization, we assessed the ability of Δ*san*_0913 to form biofilms and attach to vaginal epithelial cells. We observed no differences between COH1 and Δ*san*_0913 biofilm formation in either bacteriologic (Todd-Hewitt broth, THB) or eukaryotic (RPMI-1640) media as measured by crystal violet staining (**Fig. 2C**). Additionally, we observed no differences in VK2 adherence between the COH1 and Δ*san*_0913 strains (**Fig. 2D**). In our *in vivo* model, we found no differences in vaginal GBS burdens between COH1 and Δ*san*_0913 (**Fig. 2E**). However, when mice were treated with pHMOs as in **Fig. 1A**, Δ*san*_0913 displayed significantly higher GBS burdens at day 1 post-inoculation (*P* = 0.007) during active pHMO treatment, but this difference resolved at later time points (**Fig. 2F**). Furthermore, we performed minimum inhibitory concentration (MIC) assays of a variety of antibiotic classes, hydrogen peroxide, and dimethyl sulfoxide (DMSO). MICs were determined by a >90% reduction in OD_600_ values compared to controls. No differences in MICs between COH1 and Δ*san*_0913 were observed with any compound tested (**Supp. Table 1**).

**Figure 2.**
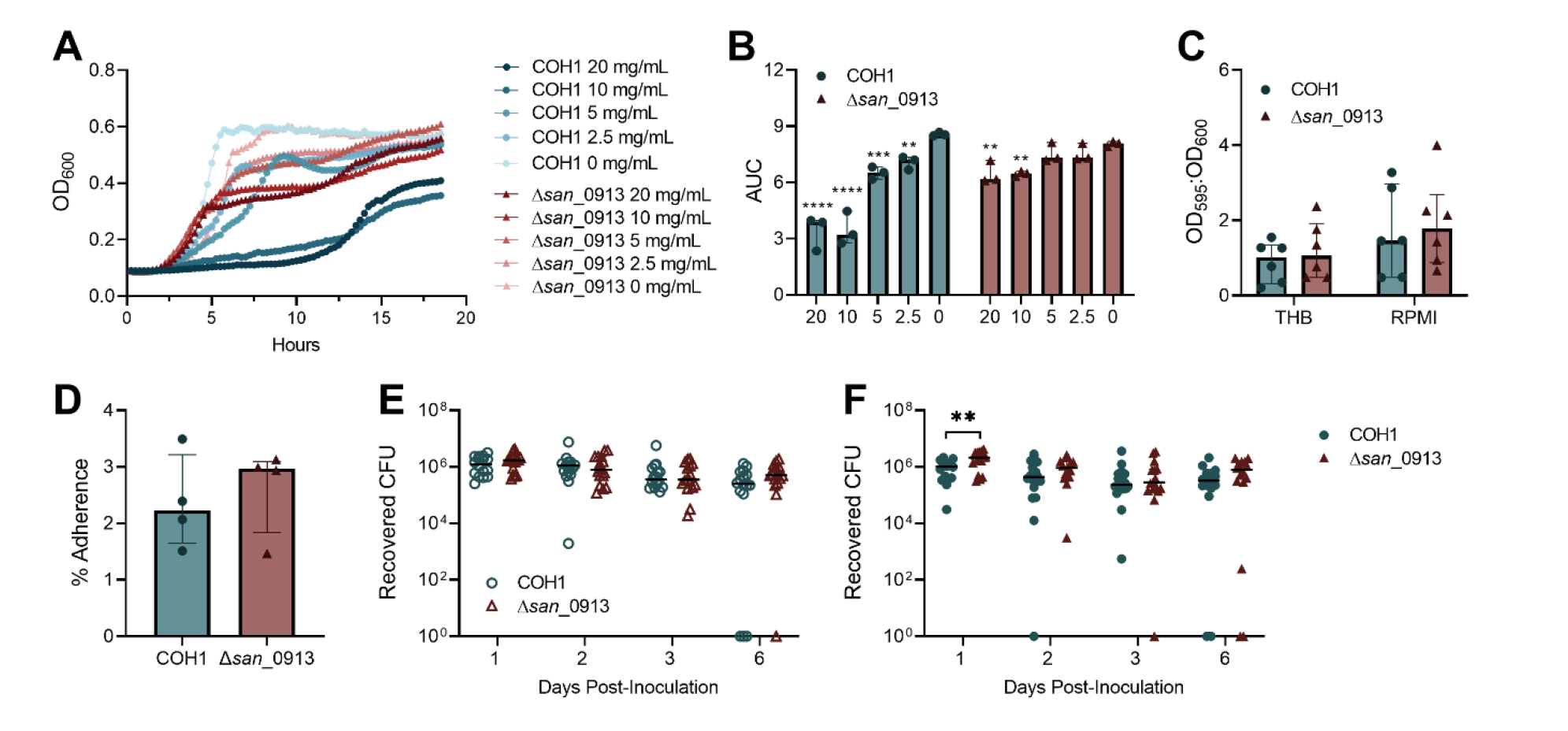
HMO resistance conferred by disruption of *san*_0913 does not alter GBS biofilms, adherence, or *in vivo* colonization in the absence of HMOs. (A) Growth curves of WT COH1 and Δ*san*_0913 in RPMI-1640 supplemented with 0, 2.5, 5, 10, or 20 mg/mL pHMOs and cultured for 18 h as measured by optical density (OD_600_). (B) Area under curve analysis of growth curves from (A). Comparisons shown are to 0 mg/mL pHMO controls. (C) Biofilm formation of COH1 and Δ*san*_0913 in THB or RPMI-1640 quantified by crystal violet staining and expressed as a ratio of crystal violet absorbance over total bacterial biomass (OD_595_:OD_600_). (D) Percent adherence of COH1 and Δ*san*_0913 to VK2 cells after 30 min of infection, MOI = 1. (E) Mice were vaginally inoculated with 1 × 10^7^ CFU of COH1 or Δ*san*_0913, and vaginally swabbed at indicated time points. Recovered GBS CFU recovered from swabs are shown. (F) Mice were inoculated as in (E) and treated with pHMOs as indicated in Fig. 1A. Recovered GBS CFU recovered from swabs are shown. Symbols represent the median of three independent experiments (A), means of three to six independent experiments (B-D) or individual mice from two combined independent experiments (*n* = 16/group, E,F). Lines indicate median values and interquartile ranges. Data were analyzed by two-way repeated measures ANOVA with Dunnett’s multiple comparisons test (B), two-way ANOVA with Sidak’s multiple comparisons test (C), Wilcoxon matched-pairs signed rank test (D), or two-stage Mann-Whitney test (E,F). **** *P* < 0.0001, *** *P* < 0.001, ** *P* < 0.01, *\* P* < 0.05. All other comparisons are not significant.

### pHMO treatment minimally impacts the endogenous murine vaginal microbiota in the presence or absence of GBS

We previously identified that GBS introduction to the murine vaginal tract causes community instability, particularly a decrease in *Staphylococcus succinus*, a dominant vaginal microbe in C57BL/6J mice (33). Because HMOs are metabolized by a variety of bacteria in the neonatal intestinal tract (34-38), and since maternal serum HMO levels correlate with specific taxa in the maternal urinary and vaginal microbiota (39), we investigated whether pHMO treatment impacted the murine vaginal microbiota in the presence or absence of GBS perturbation. Using swabs collected from the murine experiments as outlined in **Fig. 1A**, 16S rRNA amplicon sequencing was used to characterize shifts in the vaginal microbiota of Control (mock-treated, mock-infected), pHMO (treated, mock-infected), Control_GBS (mock-treated, GBS-infected), and pHMO_GBS (treated, GBS-infected) mice. The alpha diversity, as measured by Shannon’s diversity index, significantly increased in Control_GBS and pHMO_GBS groups regardless of treatment compared to controls (**Fig. 3A**). However, in the absence of GBS, alpha diversity was not impacted in the pHMO versus Control groups at any time point (**Fig. 3A**). As observed previously (33), mice that received GBS showed heightened community instability compared to mock-infected controls as measured by Bray-Curtis distance between time points. This effect was seen both in the presence (pHMO_GBS, *P* = 0.0048) and absence (Control_GBS, *P* = 0.0073) of pHMO treatment for the pairwise comparisons between days 2 and 3 (**Fig. 3B**). No impact upon community stability was observed with pHMO treatment in the absence of GBS (pHMO, **Fig. 3B**).

**Figure 3.**
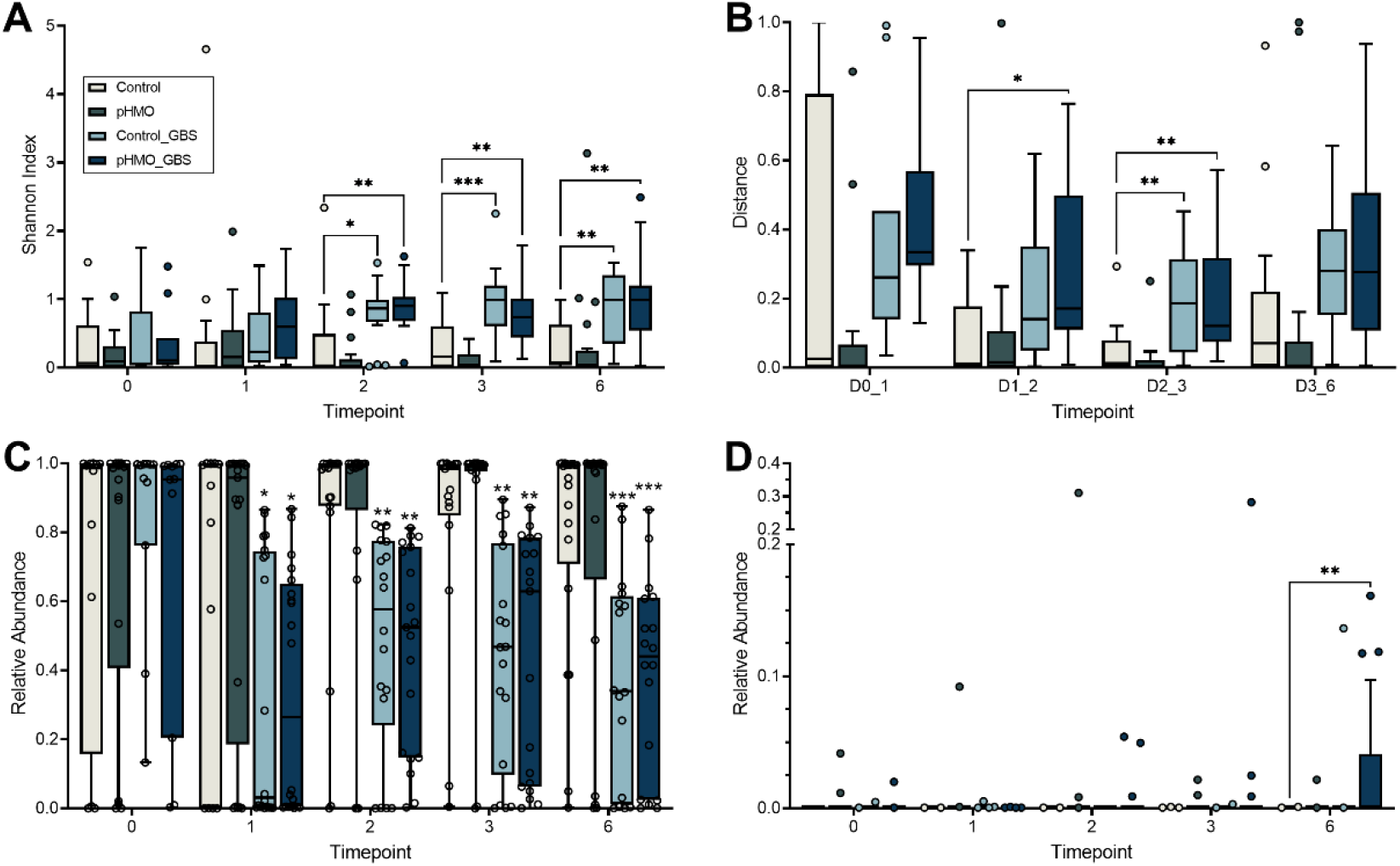
Alpha and beta diversity and differential taxa abundance as measured by 16S rRNA amplicon sequencing. Mice were mock-infected or GBS-infected and treated with pHMOs or mock-treated: Control (mock-treated, mock-infected), pHMO (treated, mock-infected), Control_GBS (mock-treated, GBS-infected), and pHMO_GBS (treated, GBS-infected) as described in Materials and Methods. (A) Shannon’s diversity index of vaginal 16S amplicon sequencing from each condition over the time course. (B) Bray-Curtis pairwise distances between subsequent time points. Relative abundance of *S. succinus* (C) and *Bacteroides spp* (D) according to treatment group over time. Displayed as Tukey’s box plot (A,B,D) and min-to-max box and whisker plots (C), *n* = 11-21/group per time point. Data were analyzed by two-way repeated measures ANOVA with Tukey’s multiple comparisons test. All comparisons shown are to the Control group. *** *P* < 0.001, ** *P* < 0.01, * *P* < 0.05. All other comparisons are not significant.

Across all four conditions, no significant differences were observed in community richness over the 6-day time course as measured by observed operational taxonomic units (OTUs, **Supp. Fig. 1A**). Mice exposed to GBS (Control_GBS and pHMO_GBS), regardless of treatment, experienced a significant drop in the relative abundance of *S. succinus* compared to Control mice starting at day 1, and this effect continued throughout the sampling period (**Fig. 3C**). No differences in the relative abundance of *Enterococcus* spp. or *Lactobacillus* spp., the two next most abundant endogenous OTUs, were observed between groups (**Supp. Fig. 1B, 1C**). ANCOM analysis (40) identified *Bacteroides* as the only significantly differentially abundant taxa across the four groups, with increased abundance in pHMO_GBS mice compared to all other groups (**Fig. 3D**).

### Murine vaginal community state types (mCSTs) display minimal differential stability upon pHMO treatment in the presence or absence of GBS

The human vaginal microbiome, and more recently the murine vaginal microbiome, are classified into community state types (CSTs) (41) and murine community state types (mCSTs) respectively (33). In humans, four of the CSTs are each dominated by different *Lactobacillus* species, and the remaining CSTs had a non-*Lactobacillus* dominant taxa or diverse array of facultative and strictly anaerobic bacteria (41). In C57BL/6J mice from Jackson Labs, the vaginal microbiome is separated into 5 mCSTs dominated by either *S. succinus, Enterococcus*, a mixture of *S. succinus*/*Enterococcus, Lactobacillus*, or a mixture of different taxa (33). In this study, we detected all five of these mCSTs by hierarchical clustering with Ward’s linkage of Euclidean distances of day 0 swab samples prior to GBS infection and/or pHMO treatment (**Supp. Fig. 2**). When analyzing the collection of samples from all four groups across all time points, we observed the emergence of three GBS-containing groups: GBS dominant (mCST VI), GBS and *S. succinus* present at similar levels (mCST IV), and *S. succinus* dominant with lower abundances of GBS or *Enterococcus* (mCST II) (**Fig. 4**).

**Figure 4.**
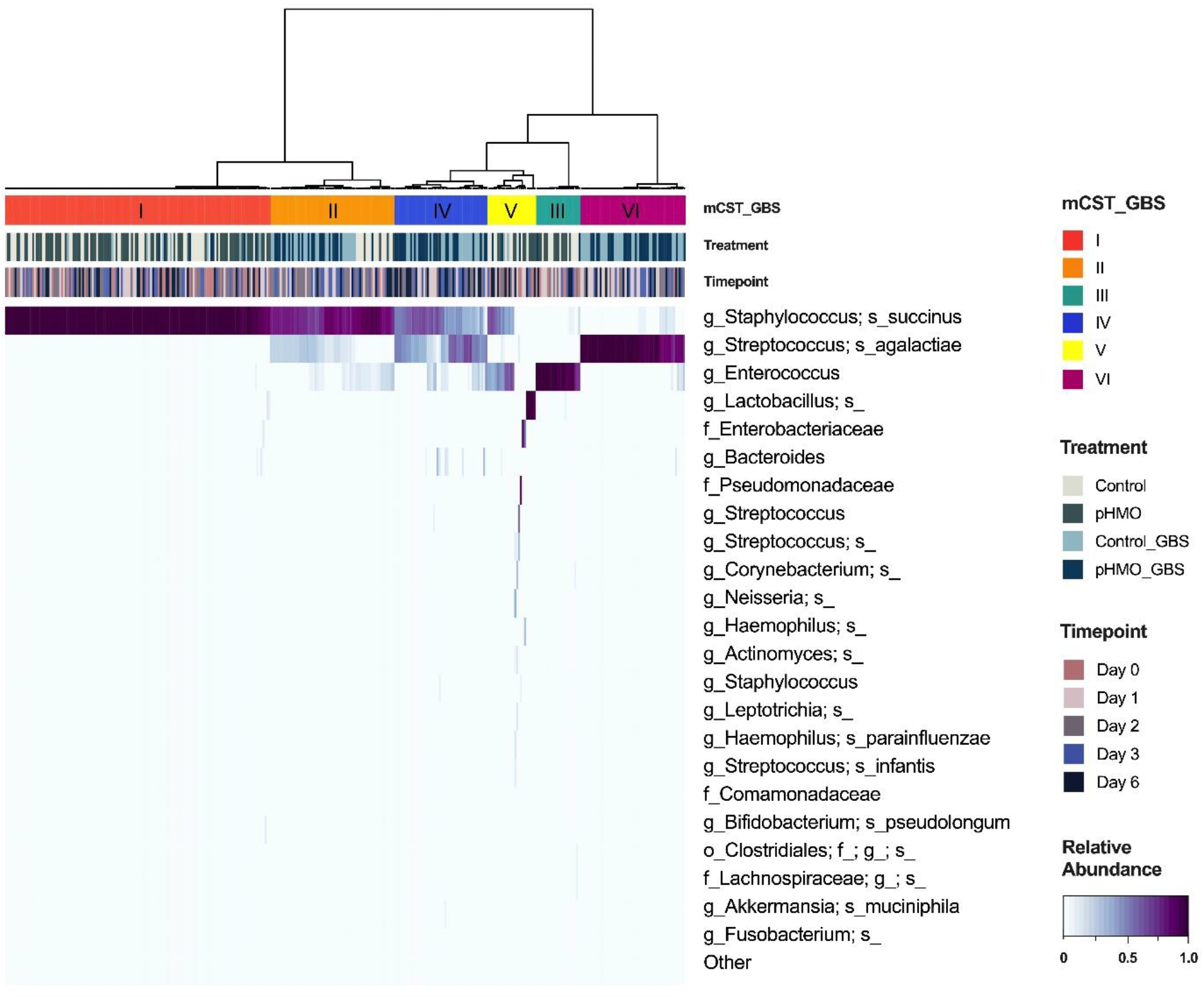
Heatmap of murine community state types across treatment groups and time points. Relative abundances of the top 23 taxa in mice across all four treatment groups as determined by 16S rRNA amplicon sequencing (*n* = 20-24 mice/group). Murine samples are hierarchically clustered by Ward’s linkage of Euclidean distances to generate mCSTs (top bar). Treatment (middle bar) and timepoint (bottom bar) per sample are displayed above the heatmap. Highest to lowest taxonomic abundances are shown by heatmap intensity corresponding to the colorbar (indicated in lower right corner) ranging from dark purple to white.

To assess if mice differentially transitioned between mCSTs across treatment groups, we tracked mCSTs in individual mice over time. Like our prior study (33), we found that mCSTs were relative unstable, with 43% of uninfected and 87% of GBS-infected mice being categorized to two or more mCSTs over the time course (**Fig. 5**). Using Bray-Curtis first distances for microbial communities within individual mice, we compared the instability between the baseline composition and the subsequent time points. Although there were no differences in longitudinal stability between Control and pHMO groups (*P*= 0.2036), Bray-Curtis first distances were higher in pHMO_GBS versus Control_GBS mice (*P=* 0.0281) (**Fig. 5**).

**Figure 5.**
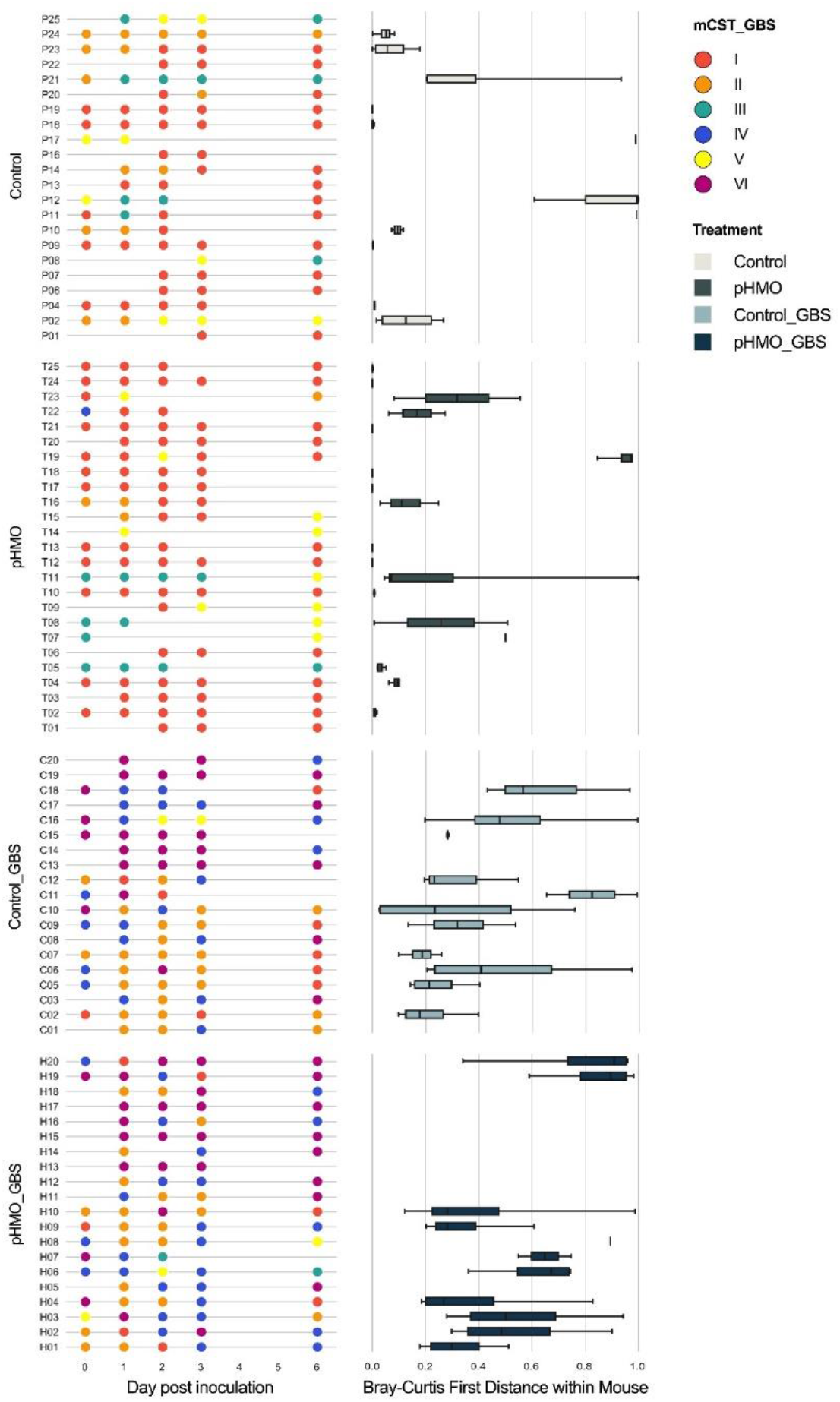
Vaginal microbiome stability over time with pHMO treatment and/or GBS infection. mCST designations for mouse cohort samples are displayed ordered by treatment group and time point (left panels). For each mouse, corresponding Bray-Curtis first distances from the day 0 time point are shown (right panels). Mice with less than two sequenced samples were excluded from analysis, and mice without a sequenced day 0 sample were excluded from the first distance analysis (*n* = 20-25/group). Data were analyzed by Mann-Whitney test.

Although mCST I (*S. succinus* dominant) was the most commonly appearing mCST in Control and pHMO groups, mCST II appeared with significantly more frequency in the Control group (*P=* 0.0404) and mCST I appeared with more frequency in the pHMO group (*P=* 0.0067) (**Fig. 6A**). No significant differences in mCST frequencies were observed between Control_GBS and pHMO_groups with mCST II, mCST IV, and mCST VI representing the most abundant mCSTs in both GBS-infected groups (**Fig. 6A**). As seen previously (33), mCST I was the most stable community state: combining all conditions and samples with successfully sequenced consecutive timepoints, 84/109 (77%) of mCST I samples were assigned mCST I at the next time point (self-transition). mCST VI (GBS dominant) was the next most stable, followed by mCST II, mCST III, mCST V, and mCST IV (**Fig. 6B**). When separated by treatment groups, we found that mCST I was more likely to self-transition in the pHMO group compared to Control group (*P* = 0.0401) whereas mCST II was more likely to self-transition in the Control group compared to the pHMO group (*P* = 0.0031) (**Fig. 6C**). In GBS-infected animals, no significant differences in mCST self-transitions were observed between Control_GBS and pHMO_GBS groups (**Fig. 6C**).

**Figure 6.**
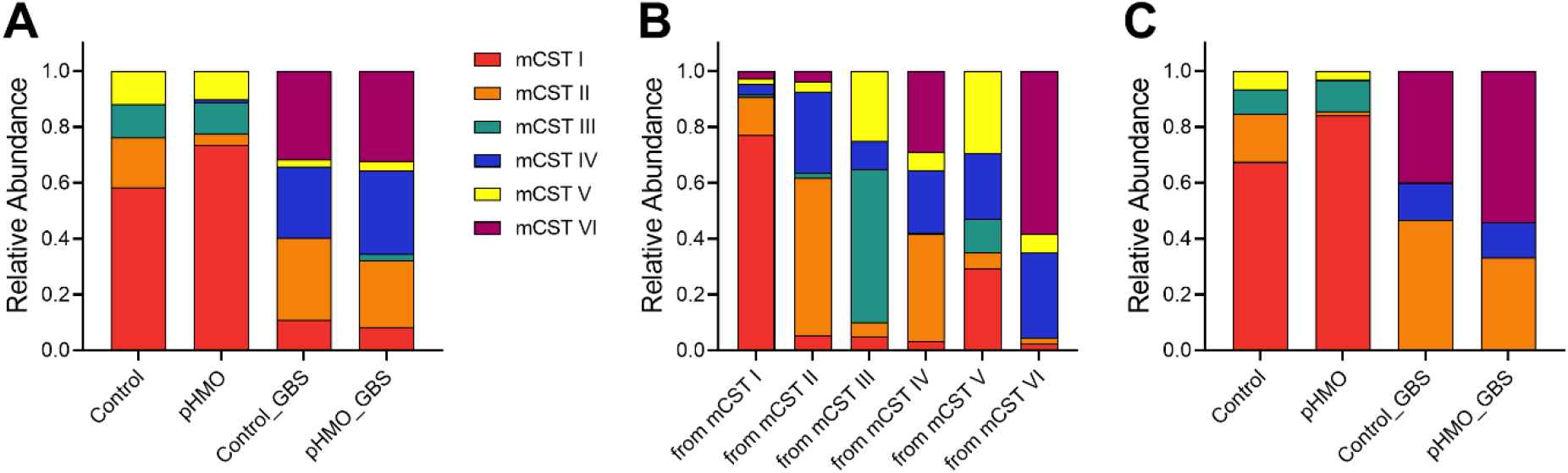
Frequency and transitions of mCSTs across treatment groups. mCST designations for mouse cohort samples were combined from all time points. (A) Frequency of mCST appearances within treatment groups. (B) Proportion of samples designated to each mCST grouped by the mCST from the previous time point. A self-transitioning mCST would be designated from a mCST to the same mCST at the next time point (e.g., from mCST I to mCST I). (C) Relative proportions of mCSTs that self-transitioned at the next time point separated by treatment group. Data were analyzed by chi square test.

## DISCUSSION

GBS remains a pervasive pathogen in pregnancy and the neonatal period. Current IAP prevention strategies have not fully abolished GBS neonatal infections and IAP is ineffective in preventing GBS infection prior to parturition. Because of the adverse effects of antibiotic exposure on the endogenous microbiota and propagation of antibiotic resistance, discovery of more targeted antimicrobial therapies to control maternal GBS carriage is important for maternal and neonatal health. Here, we apply HMOs, natural products produced by the mammary gland during pregnancy and lactation, to *in vitro* and murine models of GBS vaginal colonization. HMOs are known for simultaneous prebiotic benefits on commensal bacteria (14, 34, 38) and antimicrobial activity towards pathogens including GBS (21-24). To our knowledge, this is the first application of HMOs as a vaginal therapy *in vivo*. We propose that HMOs possess promising anti-GBS activity in this environment with minimal impact on the vaginal microbiota.

Our animal model demonstrated that pHMO treatment reduced GBS vaginal carriage, but this effect was only seen during active treatment with no sustained impact observed after treatment ceased (**Fig. 1**). This finding aligns with other murine models showing protective effects of HMOs in reducing pathogen colonization (31, 42-44). Using human vaginal epithelial cells (VK2), we observed no changes in bacterial adherence when cells were pretreated with pHMOs. This observation is distinct from work showing HMO-mediated inhibition of pathogens (31, 45-47) or enhanced attachment of beneficial bacteria (32, 48-50) at the gastrointestinal mucosa. Other studies have observed no impact of pHMO treatment on certain pathogens (51) or on pathogen colonization of other epithelial surfaces such as the bladder (52). These results suggest that prior mechanisms seen with HMOs and the gut epithelium may absent in the vaginal epithelium or with the bacterial species we tested.

There are several limitations to this HMO treatment model. First, we did not optimize dosage, timing, or length of pHMO treatment. Second, although LNT shows potent *in vitro* anti-GBS activity (21), this did not translate to an *in vivo* GBS reduction, and thus the specific HMOs responsible for GBS reduction in our animal model are currently unknown. A clinical study found that Lewis positive women, who generate certain fucosylated HMOs, display reduced GBS vaginal carriage and infant colonization at birth (27). Specifically, levels of lacto-N-difucohexaose I (LNDFHI) in breastmilk samples negatively correlated with maternal GBS colonization status and reduced GBS growth *in vitro* (27). Third, HMOs and their fermentation products have multiple known gastrointestinal epithelial and immune modulatory activities (53-56). Likewise, it is possible that HMOs can act indirectly through altering host vaginal responses to GBS, however this was not evaluated in our study. Lastly, using murine models to test whether HMOs possess potential therapeutic activity in preventing GBS neonatal transmission and adverse birth outcomes (57, 58) will be an important application of our findings.

Although the exact mechanism of anti-GBS activity by HMOs is unknown, GBS susceptibility is linked to expression of a GBS-specific putative glycosyltransferase (locus *san*_0913) thought to catalyze the addition of glucose or galactose residues to the cell surface and thus may enable incorporation of HMOs into the GBS cell wall (21). In prior work, a glycosyltransferase-deficient Δ*san*_0913 strain showed resistance to HMO inhibition (5 mg/mL) over 7 h of culture (21). In our growth analysis, we confirmed this finding extended out to 18 h (**Fig. 2**). At higher concentrations (10 and 20 mg/mL) matching physiologic concentration of HMOs in human colostrum and breastmilk (59, 60), Δ*san*_0913 growth was inhibited, but not to the same extent as wild type COH1, suggesting that this deficiency does not completely resolve anti-GBS activity of HMOs. Recent work has shown that HMOs induced multiple GBS stress responses related to cell membrane and cell wall components (26), but the role of *san*_0913 in this GBS response has not been established. While streptococcal glycosyltransferase activity has been implicated in biofilm formation and composition in *S. mutans* (61), our phenotypic analyses did not reveal any substantial deficits in the glycosyltransferase-deficient Δ*san*_0913 in terms of biofilm formation, vaginal cell adherence, or *in vivo* vaginal colonization in the absence of HMO treatment. These results may have important clinical implications for HMO therapies and emergence of spontaneous HMO-resistant GBS under selective pressure.

HMOs serve as prebiotics for beneficial microbes in the gut by promoting the establishment of *Bifidobacteria* and *Bacteroides* (37, 38, 62). Mammary HMO production begins early in pregnancy and is detected in maternal circulation in the first trimester (63). Moreover, maternal serum levels of two abundant HMOs (2’-FL and 3’-SL) positively correlate with vaginal *Gardnerella spp*. and *L. crispatus* respectively (39), providing a basis for the hypothesis that HMOs might not only shape neonatal microbiota and immunity, but also maternal vaginal microbiota. Whether HMOs have the potential to directly impact the vaginal microbiome in humans has not been determined, however, a common vaginal species, *L. gasseri*, lacks the ability to metabolize HMOs (34). Because of the well-known prebiotic effects of HMOs on the infant microbiota, we examined the impact of pHMOs on the murine vaginal microbiota in our colonization model. We found minimal pHMO-driven changes to the community composition in terms of alpha and beta diversity (**Fig. 3**). The most marked difference between groups in our model was the emergence of *Bacteroides* in mice dually inoculated with GBS and treated with pHMOs (**Fig. 3D**). While the relative abundance of *Bacteroides* remained below 5% of the entire microbial landscape in the majority of mice, 0.1-5% abundance is estimated to account for ~10^5^-10^6^ total CFU in the murine vaginal tract. In women, the vaginal microbiota postpartum shows community instability and increases in *Bifidobacterium* and *Bacteroides* (64, 65), but the mechanisms driving these changes are unknown. Whether HMOs can be detected in the human vagina during pregnancy and lactation, and whether human vaginal microbes can metabolize HMOs are important topics of future study.

There are several limitations to the interpretation of our murine vaginal microbiome data. First and foremost, the murine vaginal microbiome does not fully reflect the human vaginal microbiome in terms of species present; although there is an mCST dominated by a murine *Lactobacillus* (**Supp. Fig. 2**), it is a rare community in C57BL/6J mice (33). As a future direction, we seek to use humanized microbiota mice to assess pHMO-mediated changes to the vaginal microbiota in the presence of human vaginal bacteria, such as that done in mouse models colonized with human gastrointestinal microbiota and treated with HMOs (42, 66). In women, GBS is present at low relative abundance in the vagina (67) whereas in our mouse model, GBS becomes a dominant member of vaginal community in some mice upon introduction (**Fig. 4**). This high relative abundance may alter dynamics of GBS and other vaginal taxa distinct from human vaginal communities. Additionally, the length of HMO treatment may need to be extended to observe larger effects. Prior studies have described more pronounced HMO-mediated shifts to the gut microbiota of both conventional (44, 68) and humanized microbiota mice (66), however, the length of HMO treatment in these studies was longer than in our model (3-8 weeks vs. 3 days respectively).

By combining our prior (33) and current studies, we found that the vaginal microbiome of the C57BL/6J mice from Jackson labs is highly consistent across cohorts over several years. In both studies, we found that GBS introduction increases vaginal community instability and reduces the relative abundance of the most abundant taxa *S. succinus*. Additionally, we confirmed our prior observation that mCST I (*S. succinus*-dominant) is the most stable murine community over time. These consistencies highlight the utility of this murine model in comparing different experimental groups across cohorts and experimental variables.

In summary, we have demonstrated HMOs can reduce GBS vaginal colonization in an animal model with minimal impacts on the vaginal microbiota. There is mounting evidence that HMOs play an important role in shaping the infant gut microbiota and preventing pathogen colonization. HMO introduction to the vaginal tract may provide similar beneficial effects. These findings expand our knowledge of therapeutic applications of HMOs and support their continued development as a target for controlling GBS colonization in women.

## MATERIALS & METHODS

### Reagents, bacterial strains, and cell lines

Pooled HMOs were isolated from human milk samples collected through the human milk donation program at the University of California, San Diego, lyophilized and stored at -20° C as previously described (69). Individual HMO lacto-N-tetraose (LNT) was purchased from Dextra Laboratories. Prior to use, HMOs were resuspended in molecular grade water to a final concentration of 100 mg/mL, and subsequent dilutions were made in cell culture media (*in vitro*) or molecular grade water (*in vivo*).

Group B *Streptococcus* (GBS) strains used in this study include COH1 (ATCC BAA-1176) and isogenic Δ*san*_0913 generated previously (21). Strains were grown for at least 16 h to stationary phase at 37°C in Todd-Hewitt Broth (THB) prior to experiments with 5 μg/mL erythromycin added to Δ*san*_0913 cultures. Prior to *in vitro* and *in vivo* experiments, overnight cultures were diluted 1:10 in fresh THB, and incubated stationary at 37°C until mid-log phase (OD_600nm_ = 0.4). *Lactobacillus rhamnosus* GG (ATCC 53103) was grown for 16 h to stationary phase at 37°C without shaking in de Man, Rogosa and Sharpe (MRS) broth.

Immortalized human vaginal epithelial cells (VK2/E6E7, ATCC CRL-2616) were cultured in keratinocyte serum-free medium (KSFM) (Gibco) with 0.5 ng/mL human recombinant epidermal growth factor and 0.05 mg/mL bovine pituitary extract. Cells were cultured in a 37°C incubator with 5% CO_2_. Cells were split every 3-4 days at ~80% confluency, and 0.25% trypsin/2.21mM EDTA (Corning) were used to detach cells for passaging.

### GBS growth kinetics

For growth curves, log phase GBS cultures were diluted 1:10 in RPMI-1640 (Gibco) in 96-well microtiter plates with 20, 10, 5, or 2.5 mg/mL pHMOs or carrier control in 200 µL total volume. Wells with pHMOs and media only were also included to confirm absence of microbial contamination. Plates were incubated at 37°C and absorbance at OD_600nm_ was read every 15 min for 18 h using a BioTek Cytation 5 multi-mode plate reader.

### Biofilm assays

GBS biofilm assays were performed as described previously (70). Briefly, overnight cultures were diluted to OD_600nm_ = 0.1 in RPMI-1640 or THB and incubated at 37°C for 24 h. Media was removed, and biofilms were washed twice with PBS before drying at 55°C for 30 min. Biofilms were stained with 0.2% crystal violet for 30 min, washed with PBS three times, and destained with 80:20 ethanol:acetone mixture. Supernatant was transferred to a fresh 96-well plate and absorbance was read at OD_595nm_ using a BioTek Cytation 5 multi-mode plate reader. Values were normalized to total bacterial growth prior to washing and staining and data were expressed as a ratio of crystal violet staining to total bacterial growth (OD_595_:OD_600_).

### Minimum inhibitory concentration (MIC) assays

MICs were performed as described previously with minor adaptions (71). Mid-log phase cultures were diluted 1:100 in THB with or without H_2_O_2_, DMSO (Fisher Scientific), trimethoprim (Sigma), chloramphenicol (Fisher Scientific), and vancomycin (Sigma) at concentrations listed in **Supp. Table 1** in 100 µL total volume in 96-well microtiter plates. Plates were incubated stationary for 24 h at 37ºC. The MICs were determined by at >90% reduction in OD_600_ absorbance compared to control wells.

### Adherence assays

GBS adherence assays were performed on confluent VK2 cells in 24-well plates as described previously (72, 73). For studies using HMOs, media was replaced with KSFM containing 3mg/mL or 6 mg/mL of pHMO, LNT or vehicle control for 18 h. Cells were infected with GBS COH1, Δ*san*_0913, or *L. rhamnosus* at MOI = 1 (assuming 1 × 10^5^ VK2 cells per well). Bacteria was brought into contact with the VK2 cells by centrifuging for 1 min at 300 × *g*. After 30 min, supernatant was removed and cells washed 6X with sterile PBS. Cell layers were incubated for 5 min with 100 μL 0.25% trypsin/2.21 mM EDTA after which 400 μL of 0.025% Triton-X in PBS was added. Wells were mixed 30X to ensure detachment, and bacterial recovery was determined by plating on THB or MRS agar plates using serial dilution and counting CFUs. Data were expressed as a percentage of adherent CFUs compared to original inoculum.

### Animals

Animal experiments were approved by the UC San Diego and Baylor College of Medicine Institutional Animal Care and Use Committees (IACUC) and conducted under accepted veterinary standards. Mice were allowed to eat and drink *ad libitum*. WT C57Bl/6J female mice, originally purchased from Jackson Laboratories, aged 7 weeks, were allowed to acclimate for one week prior to experiments.

### Murine GBS vaginal colonization model

Vaginal colonization studies were conducted as described previously (74). Briefly, mice were synchronized with 0.5mg β-estradiol administered intraperitoneally (i.p.) 24 h prior to inoculation. Mice were inoculated with 10μL (1×10^7^ CFU total) of GBS COH1 or PBS as a mock control into the vaginal tract. Where applicable, mice were administered 1 mg (10 μL of 100 mg/mL) pHMOs, LNT, or vehicle control into the vaginal lumen two hours post-inoculation. Vaginal swabs were collected daily and recovered GBS (identified as pink/mauve colonies) was quantified by plating on CHROMagar StrepB (DRG International Inc.). Growth of blue colonies was considered endogenous *Enterococcus* spp. based on manufacturer protocols. Where applicable, mice received additional HMO or mock treatments on days 1 and 2 immediately following swab collection. Remaining swab samples were stored at -20° C until further use.

### Sample processing and 16S rRNA amplicon sequencing

DNA was extracted from thawed bacterial swab suspensions using the Quick-DNA Fungal/Bacterial Microprep Kit protocol (Zymo Research). The V4 regions of the 16S rRNA gene were amplified using barcoded 515F-806R primers (75), and the resulting V4 amplicons were sequenced on an Illumina MiSeq. Raw sequencing data were transferred to Qiita (76). Sequences were demultiplexed, trimmed to 150-bp reads, and denoised using Deblur through QIIME2 v2020.8 (77). Qiime2 was also used for rarefaction (1900 sequences per sample), and calculation of alpha diversity (Shannon and OTUs) and beta diversity (Bray-Curtis distance). For ANCOM (40) analysis for differentially abundant OTUs, the nonrarefied feature table was used. Taxonomic assignments used the naive bayes sklearn classifier in QIIME 2 trained on the 515F/806R region of Greengenes 13_8 99% OTUs. As many of the samples were low biomass, DNA contaminants from sequencing reagents and kits had a substantial impact on the dataset. Negative controls that went through the entire pipeline, from DNA extraction to sequencing, were used to catalog these contaminants (*Pseudomonas veronii*). Mitochondria and chloroplast 16S sequences were also removed. Output files generated through the Qiime2 pipeline were exported and analyzed with R version 3.6.1 (2019-07-05) -- “Action of the Toes” using stats, factoextra, and Phyloseq (78, 79). Data visualization was performed with ggplot2 (80) and Seaborn (81).

### Community State Type (CST) delineation

Feature tables and representative sequences generated from three individual studies were merged and used to generate a taxonomy file. Two more studies from our prior work (33) were downloaded from EBI accession number PRJEB25733 in addition to the current study (EBI accession XXXX) for **Supp. Fig. 2** depicting the Baseline CSTs. To assign mCSTs and create heatmaps, hierarchical clustering was performed using the R package stats (79) on the rarefied feature table with Ward’s linkage of Euclidean distances. The optimum number of clusters (5 mCSTs) was determined using wss and silhouette (kmeans) based on the dendrogram. For EBI accession number XXXX (this study) alone, including all experimental conditions and time points, we added an additional GBS-dominant mCST as modeled by (33). For within-mouse assessment of instability and mCST transitioning, samples with only one time point collected were excluded. Samples that did not successfully sequence at the baseline (Day 0) time point were excluded from Bray-Curtis first distances analysis.

## Data availability

Sequencing Data used in this study is available in EBI under the accession number XXXX, and code is accessible at GitHub under project “XXXX”.

## Statistics

All data were collected from at least three biological replicates performed in at least technical duplicate as part of at least two independent experiments. When biological replicates were not available (e.g. immortalized cell lines and bacteria only assays), experiments were performed independently at least 3 times. Mean value from technical replicates were used for statistical analyses, with independent experiment values or biological replicates represented in graphs with mean, median with interquartile ranges, or box and whisker plots with Tukey’s as indicated in figure legends. All data sets were subjected to D’Agostino & Pearson normality test to determine whether values displayed Gaussian distribution before selecting the appropriate parametric or non-parametric analyses. In the instances where *in vitro* and *in vivo* experimental n were too small to determine normality, data were assumed non-parametric. GBS vaginal colonization burdens were assessed by Kruskal Wallis with Dunn’s multiple comparisons test or two-stage Mann-Whitney test as indicated in figure legends. GBS adherence to VK2 cells was assessed by or two-way ANOVA with Dunnett’s multiple comparisons test or Wilcoxon matched-pairs signed rank test as indicated in figure legends. GBS growth (area under curve) and biofilm formation was compared by two-way repeated measures ANOVA with Dunnett’s multiple comparisons test and two-way ANOVA with Sidak’s multiple comparisons test respectively. Data from 16S rRNA amplicon sequencing was analyzed by two-way ANOVA with Tukey’s comparison. Bray-Curtis first distances were analyzed by Mann-Whitney test. mCST transition frequencies were compared by chi square test. Statistical analyses were performed using GraphPad Prism, version 9.2.0 (GraphPad Software Inc., La Jolla, CA, USA). *P* values < 0.05 were considered statistically significant.

## AUTHOR CONTRIBUTIONS

KAP, LB, and VN conceived and designed experiments. KAP, MEM, SO, AV, PB, JZ, and DM performed experiments. KAP, MEM, and AV analyzed and interpreted results. MEM and KAP drafted the manuscript. All authors contributed the discussion/manuscript edits.

## ACKNOWLEDGEMENTS

MM and JZ were supported by an NIH T32 award (T32GM136554). KP was supported by postdoctoral fellowships from the Hartwell Foundation and the University of California Chancellor’s Postdoctoral Fellowship Program, and a Research Scholar Award from the American Urological Association. Studies were supported by the Caroline Wiess Law Fund for Research in Molecular Medicine at Baylor College of Medicine, the Burroughs Wellcome Fund Next Gen Pregnancy Initiative (NGP10103), and by a Seed grant made available through the UC San Diego Larsson-Rosenquist Foundation Mother-Milk-Infant Center of Research Excellence. The support of the Family Larsson-Rosenquist Foundation is gratefully acknowledged. The funders had no role in study design, data collection and interpretation, or the decision to submit the work for publication.

