## Supplemental Material for "Human milk oligosaccharides reduce murine group B *Streptococcus* vaginal colonization with minimal impact on the vaginal microbiota"

### Supplemental Tables:

| | **WT COH1** (CI 95%) | **Δ*san*_0913** (CI 95%) | Concentrations tested |
|---|---|---|---|
| Chloramphenicol | **1.25** (1.25-2.5) | **2.5** (1.25-2.5) | 0.625 – 40 µg/mL |
| DMSO | **20** (20) | **20** (10-20) | 0.31 – 80% (v/v) |
| $H_2O_2$ | **0.0093** (0.0093) | **0.0093** (0.0093) | 0.0047 – 0.3% (v/v) |
| Trimethoprim | **0.3125** (0.125-5) | **0.15625** (0.125-0.25) | 0.0078 – 20 mg/mL |
| Vancomycin | **0.125** (0.125-1) | **0.125** (0.125-1) | 0.031 – 2 µg/mL |

**Supplemental Table 1. Minimum inhibitory concentrations for WT COH1 and Δ*san*_0913.** All experiments were done in at least three independent experiments in technical duplicate. Data were analyzed by Wilcoxon rank test with median values and confidence interval (CI 95%) calculated from the MICs of each independent experiment.

### Supplemental Figures:

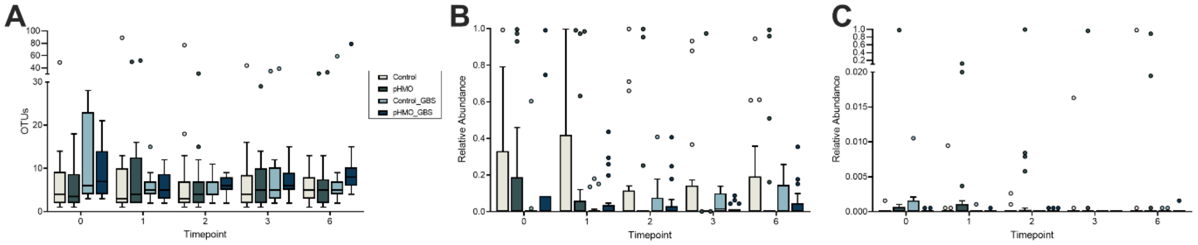


**Supplemental Figure 1. Richness and relative abundances of vaginal 16S samples.** Mice were mock-infected or GBS-infected and treated with pHMOs or mock-treatment: Control (mock-treated, mock-infected), pHMO (treated, mock-infected), Control_GBS (mock-treated, GBS-infected), and pHMO_GBS (treated, GBS-infected) as described in Materials and Methods. (A) Observed operational taxonomic units (OTUs) in mouse samples across the four experimental conditions over time. Relative abundances of (B) *Enterococcus spp.* and (C) *Lactobacillus spp.* over time. All graphs are displayed as Tukey's box plots. Data was analyzed by two-way repeated measures ANOVA with Tukey's multiple comparisons test. All comparisons are not significant.

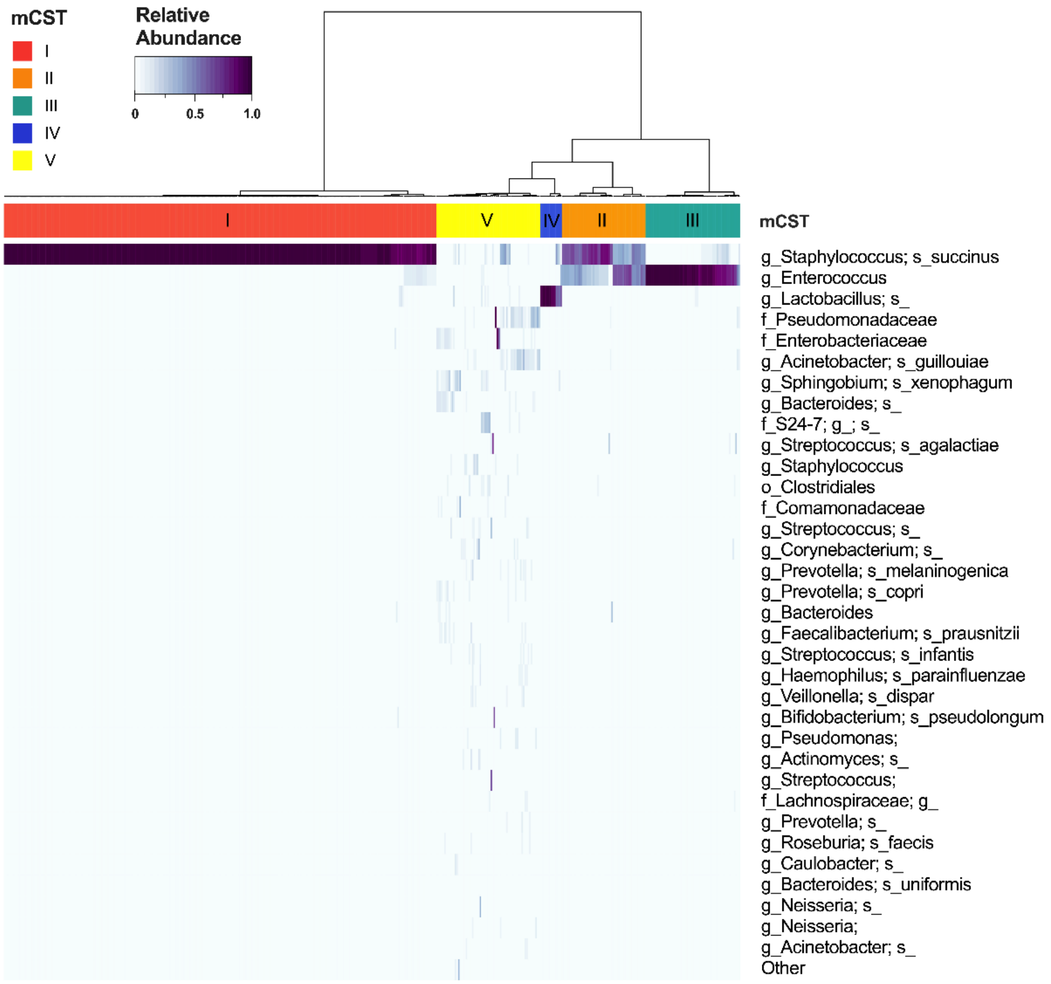


**Supplemental Figure 2. Baseline mCST clustering of C57BL/6J mouse samples from this study combined with our prior study.** Vaginal swab samples taken prior to treatment (day 0) or from mock controls (uninfected, untreated) were hierarchically clustered by Ward's linkage of Euclidean distances to generate mCSTs (top bar). The relative abundances of the top 34 taxa are displayed in a heatmap where the highest to lowest taxonomic abundances correspond to the colorbar (indicated in upper left corner) ranging from dark purple to white. Data were combined from this current study and data deposited at EBI under the accession number PRJEB25733 (Vrbanac et al, 2018, BCM Microbiology).
